# A Clarifying Perspective on Bacterial Pseudo-Receiver Domains

**DOI:** 10.1101/2025.06.23.661182

**Authors:** Robert B. Bourret, Emily N. Kennedy, Rita Tamayo, Clay A. Foster

**Affiliations:** Department of Microbiology and Immunology, University of North Carolina, Chapel Hill, North Carolina, USA; Department of Pediatrics, Section Hematology/Oncology, University of Oklahoma Health Sciences Center, Oklahoma City, Oklahoma, USA

**Author notes:** Address correspondence to Robert B. Bourret.

**Keywords:** pseudo-receiver domains, receiver domains, aspartate-less receiver domains, atypical response regulators, two-component regulatory systems

## Abstract

Two-component regulatory systems typically consist of a sensor kinase and a response regulator. All response regulators contain a receiver domain; most also contain an output domain. Response regulator activity is controlled by the phosphorylation state of the receiver. Receivers contain five conserved active site residues that catalyze phosphorylation and dephosphorylation reactions. Some protein domains identified computationally as receivers (PF00072) lack one or more of the five key conserved residues and are termed pseudoreceivers (PsRs). Because receivers are among the most abundant protein domains in nature, PsRs are also common. PsRs are especially common in plants, where they control circadian rhythms. Little is known about PsRs in fungi and archaea. Here, we focused on bacterial PsRs.

We created representative datasets of 9,153 PsR and 143,116 true receiver domain sequences from bacteria. Comparison of amino acid composition of PsR and true receiver domains at each position showed (i) many differences at positions known to be important for phosphorylation-mediated signaling in true receiver domains, consistent with diminished importance in PsRs; and (ii) greatest differences between PsR and true receiver domains in the β3α3 and β4α4 loops, potentially highlighting functionally important regions of PsRs. We also performed covariation analyses of PsR and true receiver domains, which suggested six networks of linked residues that may be important for PsR function. Our analyses lay the foundation for rational experimental approaches to investigate molecular mechanisms of signaling by bacterial PsRs.

## Nomenclature of two-component regulatory systems

Two-component regulatory systems (TCSs) mediate signal transduction in bacteria, archaea, eukaryotic microorganisms, and plants (1–7). In their simplest form, TCSs are composed of two proteins, a sensor (or histidine) kinase and a response regulator. The sensor kinase detects environmental stimuli, which regulate autophosphorylation using ATP. Thus, input information is encoded as a phosphoryl group on a conserved histidine residue, most commonly in a HisKA or HisKA_3 domain. The phosphoryl group is transferred to a conserved aspartate residue in the receiver domain of the partner response regulator. The receiver domain exists in equilibria between inactive and active conformations. The conformation of the receiver in turn controls the output function of the response regulator to implement adaptive responses to stimuli, typically via regulation of transcription. Phosphorylation stabilizes active receiver domain conformations and thus regulates output. Some TCSs contain an additional histidine-containing phosphotransfer (Hpt) domain to form a reversible multistep phosphorelay.

Receiver domains have a (βα)5 structure with a central five-stranded parallel beta sheet surrounded by five alpha helices (4, 8) (**Fig. 1A**). An active site formed by five conserved residues located on loops at the C-terminal ends of adjacent beta-strands catalyzes response regulator phosphorylation and dephosphorylation reactions. The Asp phosphorylation site (termed D) is in the center and surrounded by two acidic residues (termed DD), a Ser/Thr (termed T), and a Lys (termed K). T and K each bind to one of the three phosphoryl group oxygen atoms. The third oxygen is coordinated by a divalent cation, typically Mg^2+^, bound by DD. Although there appears to be a consensus on categorizing receiver domains that deviate from the norm, the nomenclature used to describe them is not consistently applied in the literature. To be clear, we define receiver domains that lack any one of the five conserved residues as “pseudo-receiver” (PsR) domains. We call response regulators with PsR domains “atypical response regulators” (ARR). A subset of PsRs lack the D site of phosphorylation and are termed “aspartate-less receiver” or ALR domains. Readers should beware of misleading terms that propagate through the PsR literature - for example saying the VemR response regulator is an ARR because it consists of a single receiver domain (9), the SsoR response regulator is an ARR because it is active in the absence of phosphorylation (10), or that NnaR is an orphan response regulator (11) because NnaR lacks a receiver domain (12).

**FIG 1.**
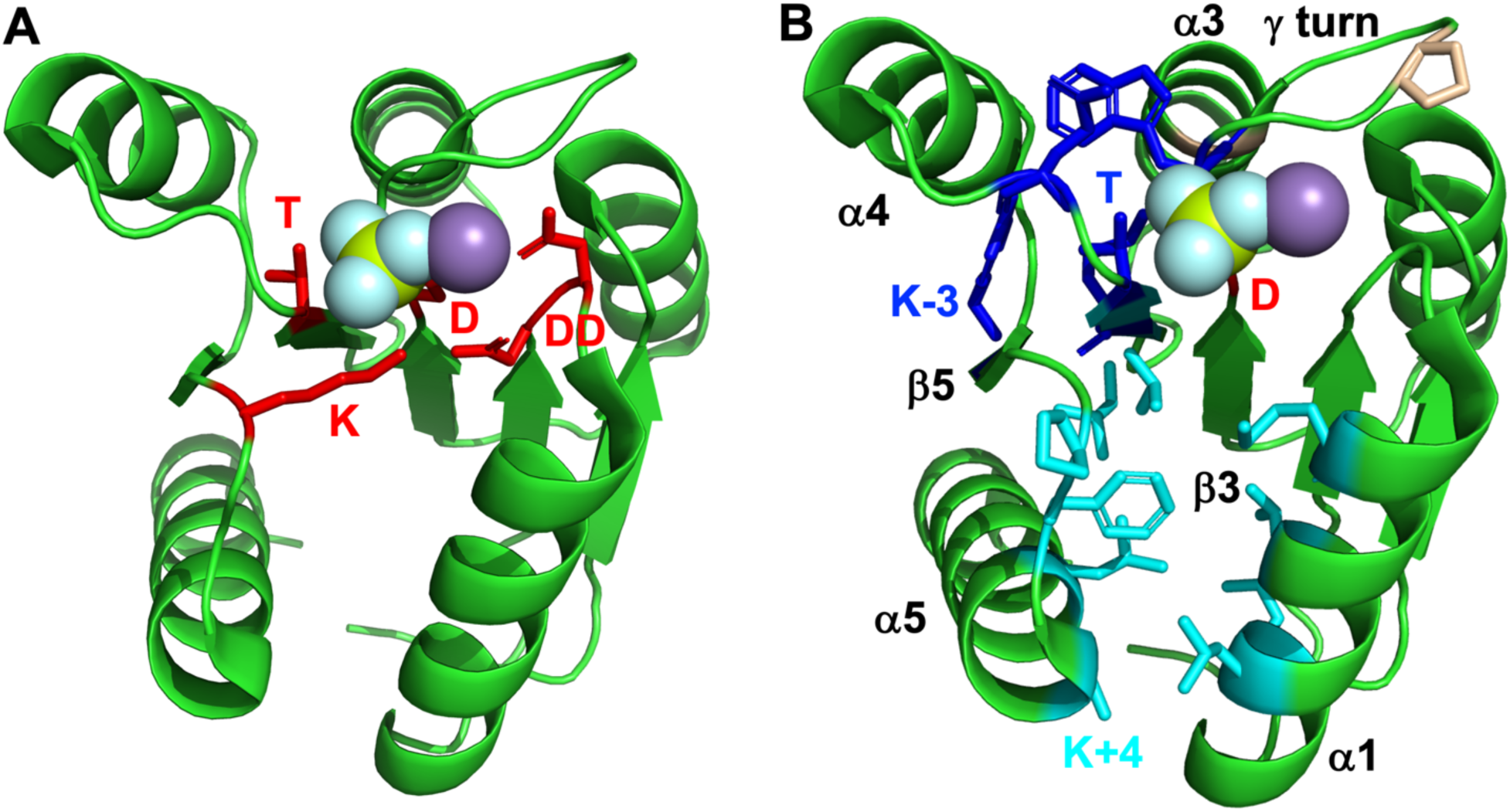
Receiver domain structure and key positions. The structure is *E. coli* CheY [PDB ID 1fqw (110)]. The purple sphere is Mg^2+^. The green and cyan spheres are the phosphoryl group analog BeF3^-^. (A) The sidechains of five conserved receiver domain residues that catalyze phosphorylation and dephosphorylation reactions are shown in red. (B) Two known allosteric networks extending between the phosphorylation site (red) and the α4β5α5 or α1α5 surfaces are shown in blue (4, 43) and cyan (44) sidechains respectively. The blue network includes T and an aromatic residue at K-3 and is commonly termed “Y-T” coupling. The γ turn between β3 and α3 is indicated, along with the relatively conserved Pro at D+4 and Gly at D+8 (tan sidechains).

## Frequency and significance of receiver and PsR domains

There are ∼14,000 different known protein domains (13). Receiver domains are in the top 0.1% of natural abundance. Protein databases such as InterPro (13) and SMART (14) use Hidden Markov Models that recognize patterns associated with the described structural features to identify amino acid sequences predicted to encode receiver domains. We previously inferred that ∼13% of domains identified as receiver domains in prokaryotic protein databases are actually PsRs (15). A more direct analysis for this study found ∼6% PsRs among a representative set of bacterial receiver domains (see Supplemental Material). Maule *et al.* found that ∼4% of domains identified as receivers across all branches of life are ALR domains, a subset of PsRs (16). About 10-15% of archaeal receiver domains (16) are ALRs, yet we are not aware of any published investigations of the function of archaeal PsRs. About 5% of fungal receiver domains are PsRs, with most instances being in Rim15p proteins (17). Rim15p’s from Ascomycota generally contain PsRs in which the D position is replaced with Glu, whereas Rim15p’s from non-Ascomycota fungal species mostly contain true receiver domains. To the best of our knowledge, the only published characterization of a fungal PsR involves a frameshift mutation that truncates the C-terminal PsR domain of *Saccharomyces cerevisiae* Rim15p and alters a phenotype, suggesting the PsR domain is important for function (18). In contrast, PsRs represent one of four major classes of response regulators in plants (19, 20). Plant pseudo-response regulators (PRRs) that contain PsRs are key components of the circadian clock and function via oscillating cycles of protein phosphorylation (not on D, which is typically replaced by Glu in plant PRRs), protein/protein interactions, protein degradation, and DNA binding to repress gene expression (21–23). Thus, PsRs are abundant in nature, yet remarkably little is known about how PsRs function outside of plants.

The InterPro database contains ∼2 million receiver domains (Pfam designation PF00072), several percent of which are PsRs. Although most experimentally characterized PsRs are classified as receiver domains by domain recognition software, a significant fraction of domains labelled as PsRs in the literature instead come from PF06490, PF21155, PF21194, PF21695, PF21714, and PF22368, far smaller Pfam groupings with tens to thousands of InterPro entries apiece. Although the six minor PsR domain types all have the same (βα)5 topology as receiver domains, the Hidden Markov Model signatures that define each do not include any of the five conserved active site residues found in receiver domains (except for D in PF21695), consistent with regulatory mechanisms that do not involve Asp phosphorylation.

## Toward conceptual clarity regarding PsRs

We intentionally define and distinguish four somewhat related topics that are often entangled in the existing literature on bacterial PsRs (16, 24–26):

i. PsRs/ARRs/ALRs, which lack one or more of the five conserved active site residues that catalyze receiver domain phosphorylation and dephosphorylation chemistry. Key unknown aspects of PsR-containing proteins are the mechanism(s) of activation and inactivation, which intuitively leads to the other three topics.
ii. Non-canonical response regulators that use phosphorylation-independent regulatory mechanisms such as allosteric coupling activation by sensor kinases (27) in addition to Asp phosphorylation-dependent mechanisms. It is conceivable, although not proven, that PsRs might employ some of the same phosphorylation-independent mechanisms as non-canonical response regulators.
iii. Canonical response regulators that use Asp phosphorylation-dependent regulation but exhibit activity in both non-phosphorylated and phosphorylated states, for example (10, 26, 28, 29). It is natural to look to such examples for clues as to how PsRs might be active without phosphorylation. However, there is no obvious reason to think that non-phosphorylated response regulators should not have activity. Instead of using two states as an on/off switch, there could be situations in which it makes sense to utilize two states to achieve either/or outcomes instead.
iv. “Orphan” response regulators without an obviously associated sensor kinase (25, 30). Genes encoding partner sensor kinases and response regulators are often found in the same operon to allow positive autoregulation and implementation of appropriate responses when the matching environmental stimulus is detected (31).

Orphan response regulator genes raise the question of whether a partner sensor kinase is encoded elsewhere or if the orphan functions independently of phosphorylation, as do PsRs. As a result, although most orphan response regulator genes encode typical rather than atypical RRs (30), the set of experimentally characterized ARRs is greatly enriched for orphans. In fact, of the 20 bacterial ARRs specifically described in this study, only CmrT and FrzS are not orphans.

Untangling the four concepts described above potentially provides an opportunity to clearly examine the PsR field from a new perspective. We will focus here exclusively on the first topic, PsRs, in bacteria. We incorporate structural analyses and literature reports of PsRs from the six minor domain families, but for practical reasons restrict all PsR sequence analyses to the predominant PF00072 class.

## A broader perspective on pseudo-TCS enzymes

Collins and Childers recently expanded the field to consider pseudo-histidine kinases (24). Histidine kinases typically support multiple functions, including stimulus detection, ATP binding, autophosphorylation, serving as a phosphodonor for response regulators, and acting as a phosphatase toward partner response regulators. Pseudo-histidine kinases lack one or more such functions but nevertheless can be integrated into TCS signal transduction networks. Consideration of partially defective TCS proteins potentially represents the start of a coherent logical framework to understand PsRs. We analyzed sequences of bacterial HisKA, HisKA_3, and Hpt domains from a Representative Proteome 75 (32) dataset (details in Supplemental Material) and found the frequency of missing His phosphorylation sites to be 0.9% (722 of 76,471 sampled) for HisKA domains, 0.3% (56 of 16,025) for HisKA_3 domains, and 0.9% (90 of 10,312) for Hpt domains. The frequency of missing Asp phosphorylation sites for “receiver” (i.e. PF00072) domains was 2.0% (3,111 of 152,269 sampled) (16). The higher frequency of missing phosphorylation sites in domains identified as receivers compared to domains identified as HisKA, HisKA_3, or Hpt is consistent with different roles of Asp and His phosphorylation. Asp phosphorylation is used to stabilize specific receiver domain conformations, which presumably can also be achieved by other means. In contrast, the primary known role of His phosphorylation sites in TCS proteins is to serve as a phosphodonor (HisKA, HisKA_3, Hpt) or phosphoacceptor (Hpt only) for receiver domains. The His phosphorylation site cannot be bypassed for phosphotransfer reactions.

The conserved His residue is not essential for the phosphatase activity mediated by HisKA or HisKA_3 domains (33–35), but contributes to the HisKA phosphatase reaction (36). In contrast, the conserved His does not appear to participate in phosphatase activity mediated by HisKA_3 domains (37). Therefore, if phosphatase activity were a primary function of pseudo-histidine kinases, then we might expect to observe a higher frequency of HisKA_3 than HisKA domains lacking the conserved His, but the opposite was observed.

## Differences in amino acid composition at key positions of PsR and receiver domains

With about 2M “receiver” domain (PF00072) sequences in InterPro, there are potentially tens or hundreds of thousands of available PsR domain sequences. We sorted PsR domains from true receiver domains as described in Supplemental Material to create a representative database (**Dataset S1**) of 9,153 bacterial PsR domain sequences. To gain insight into mechanisms used by PsRs, we analyzed the resulting multiple sequence alignment in three ways:

First, we examined the distribution of amino acids (i.e., composition) at various positions known to be important for receiver domain function (**Datasets S3, S4**). By definition, the abundance of each of the five conserved active site amino acids (**Fig. 1A**) was lower in PsRs than receivers (**Table S1A**). Recall the DD residues in receiver domain active sites are involved in metal ion binding (38). DD1 (the N-terminal residue in the DD pair) binds the metal ion via a water molecule whereas DD2 directly binds the metal ion. The amino acid composition at DD2 is more diverse between PsRs and receivers than DD1, which was about 20% non-acidic residues in PsRs. The D, T, and K positions, all of which interact directly with the receiver domain phosphoryl group, each decreased from ∼100% conservation in receiver domains to ∼65% abundance in PsRs, consistent with reduced involvement of phosphorylation compared to receiver domains. Variable residues are named relative to the landmarks of conserved positions. For example, T+1 is the position one residue to the C-terminal side of the conserved T. The amino acids at position T+1 of receiver domains generally have small side chains to allow steric access to the phosphorylation site (39, 40). The abundance of Ala and Gly at T+1 declined from 74% in receiver domains to 43% in PsRs (**Table S1B**), again consistent with a lack of a phosphorylation-dependent function in PsRs. Similarly, positions D+2 and T+2, which affect receiver domain phosphorylation and dephosphorylation reaction kinetics by interacting with attacking or leaving groups (15, 41, 42), exhibited substantially different amino acid composition between receivers and PsRs (**Table S1B**).

Receiver domains typically contain a hairpin γ-turn in the β3α3 loop, which is facilitated by a Pro at D+4 (40). Position D+8 near the N-terminal end of α3 is typically a Gly due to steric constraints on the sidechain (40). The abundance of the preferred amino acids at both positions (tan residues in **Fig. 1B**) were substantially reduced in PsRs compared to receiver domains (**Table S1C**), suggesting altered structural features as discussed in a later section.

Many receiver domains contain two well-characterized allosteric pathways that connect the site of phosphorylation with surfaces on the receiver that can form regulated protein/protein interactions. One is termed “Y-T coupling” and involves coordinated conformational changes between an aromatic residue at the K-3 position in β5 and the conserved T that link the phosphorylation site to the α4β5α5 surface (blue residues in **Fig. 1B**) (4, 43). The abundance of Tyr/Phe and Thr/Ser residues at the “Y-T coupling” positions declined from 86% and 100% respectively in receiver domains to 65% and 70% respectively in PsRs (**Table S1D**). Consequently, only 47% of PsRs retain the amino acids at both positions T and K-3 needed to support “Y-T” coupling (**Dataset S5**), compared to 86% of receivers. Similarly, the composition of nearby positions D+1, T-2, and T+2 implicated in Y-T coupling (43) were substantially different between receiver and PsR domains, again consistent with reduced importance of phosphorylation mediated signaling in PsRs.

A second allosteric pathway connects the site of phosphorylation to the α1α5 surface (cyan residues in **Fig. 1B**) (44). The amino acid composition at most positions in this pathway were similar between PsR and receiver domains (**Table S1E**). However, composition of the physically adjacent DD+4, T-1, and K+2 positions differed the most, a point to which we will return.

## Overall differences in amino acid abundance in PsR and receiver domains

Instead of restricting our view to positions known to be important in receiver domains, we also used the multiple sequence alignment data to examine in an unbiased manner the differences in abundance of each amino acid between receiver and PsR domains at all positions (**Fig. 2**). The heatmap is scaled to show positions at which amino acids differ in absolute abundance by up to 20%. Several features of the heat map are striking:

**FIG 2.**
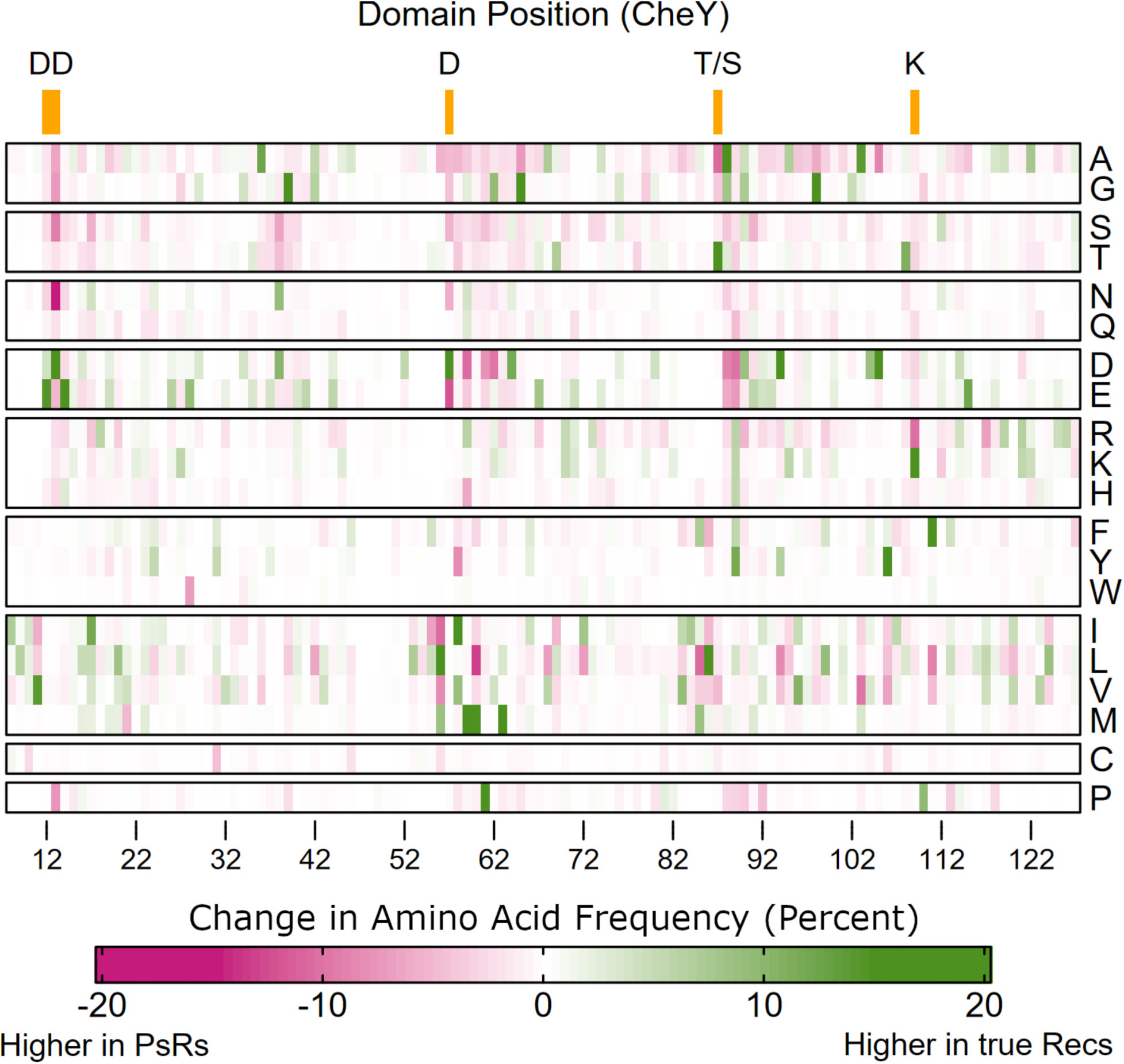
Heatmap of amino acid composition differences between receiver and PsR domains. Multiple sequences alignments of bacterial PsR and receiver domains (**Datasets S1 and S2** respectively) were constructed as described in Supplemental Material and used to calculate the percent abundance of each amino acid at every position (**Datasets S3 and S4**), with numbering according to the *E. coli* CheY sequence. PsR abundances were subtracted from receiver abundances and the differences used to generate a heat map, with greens indicating higher abundance in receiver domains and reds indicating higher abundance in PsR domains.

There were many more positions at which a particular amino acid was substantially more abundant in receiver than PsR domains (dark green bars in **Fig. 2**) than vice versa (dark magenta bars). A simple interpretation is that green bars represent positions in which particular amino acids are preferred in receiver domains and the selective pressures to maintain those preferences has been lost in PsRs due to mechanistic/functional differences between PsRs and receivers, such as the role of phosphorylation.

The positions with the strongest signal in PsRs (magenta in **Fig. 2**) generally corresponded to conserved (or nearby) positions in receiver domains, so the abundance of an amino acid changed from 0% in receivers to a nonzero value in PsRs. For example, position DD2 was commonly replaced by Asn, position D was commonly replaced by Glu, position T was commonly replaced by Ala, and position K was commonly replaced by Arg. These changes are all evident in **Table S1**. The overall dearth of positions at which particular amino acids are favored in PsRs is consistent with the possibility that there are few or no functionally important amino acids shared across all PsRs. This interpretation does not exclude the possibility that there may be multiple types of PsRs, each with a distinct set of functionally important (i.e., conserved) residues, therefore confounding a straightforward analysis of amino acid composition.

The biggest clusters of amino acid composition differences were in the β3α3 (including positions D, D+1, D+2, D+3, D+4 and D+5) and β4α4 (including positions T, T+1 and T+2) loops, where there appeared to be an enrichment of negatively charged amino acids in PsRs compared to receiver domains (and corresponding overall reductions in Pro residues at position D+4 and Ala/Gly residues at position T+1).

Covariation and structural data described below support the identification of these two loop regions as primary differences between receivers and PsRs.

## Covariation analysis of PsR domain sequences

Our third use of the PsR multiple sequence alignment was to perform a corrected mutual information analysis as described in **Supplemental Materials**. Numerous mathematical methods have been created to explore the relationship between all pairs of positions in a multiple sequence alignment [reviewed in (45)], here collectively termed “covariation analyses”. The methods have different strengths and weaknesses. Covariation analysis can suggest functionally important residues and at a minimum is a useful tool to generate experimentally testable hypotheses. The particular method (46) we used here successfully predicted functionally important residues for the model receiver domain CheY (47). The top ∼9% of pairwise covariation scores in PsR domains were statistically significant, leading to several striking observations:

In addition to the five conserved active site residues that catalyze receiver domain autophosphorylation and autodephosphorylation reactions, some variable positions affect reaction kinetics. Positions D+2, T+1, and T+2 have direct effects due to their close proximity to the phosphorylation site (15, 39, 41, 42), whereas K+1 and K+2 act allosterically by affecting conformational equilibria (47). Accordingly, pairwise covariation linkages between positions D+2/T+2, D+2/T+1, T+1/K+1, and K+1/K+2 are in the top 1.0% of all connections in receiver domains (47). In PsRs, K+1/K+2 exhibited the third strongest linkage of any pair (top 0.05%) and a strong D+2/T+2 linkage was maintained. In contrast, the D+2/T+1 connection weakened (top 2.5%) and the interaction between T+1/K+1 was no longer statistically significant. Maintained covariation of K+1/K+2 [affect conformation (47)] and reduced covariation with T+1 [controls access to phosphorylation site (39)] are consistent with the idea that PsRs may place greater emphasis on conformation than on phosphorylation-mediated changes.

Predominant networks of pairwise covariation interactions can reveal regions of PsR domains that are functionally and/or structurally important. We focus on core (isolated) networks defined by covariation scores in the top 1% and comment on extended networks formed by adding connections with covariation scores that rank between the first and second percentile. Corresponding pairwise mutual information interactions were input into Cytoscape (48) and extended networks were constructed using the clusterMaker2 plugin (49) by applying a best neighbor filter (using only first-order neighbors) for manual analysis. The following six core networks were observed (**Fig. 3**) (minimum residue count of three; details in **Table S2**):

**FIG 3.**
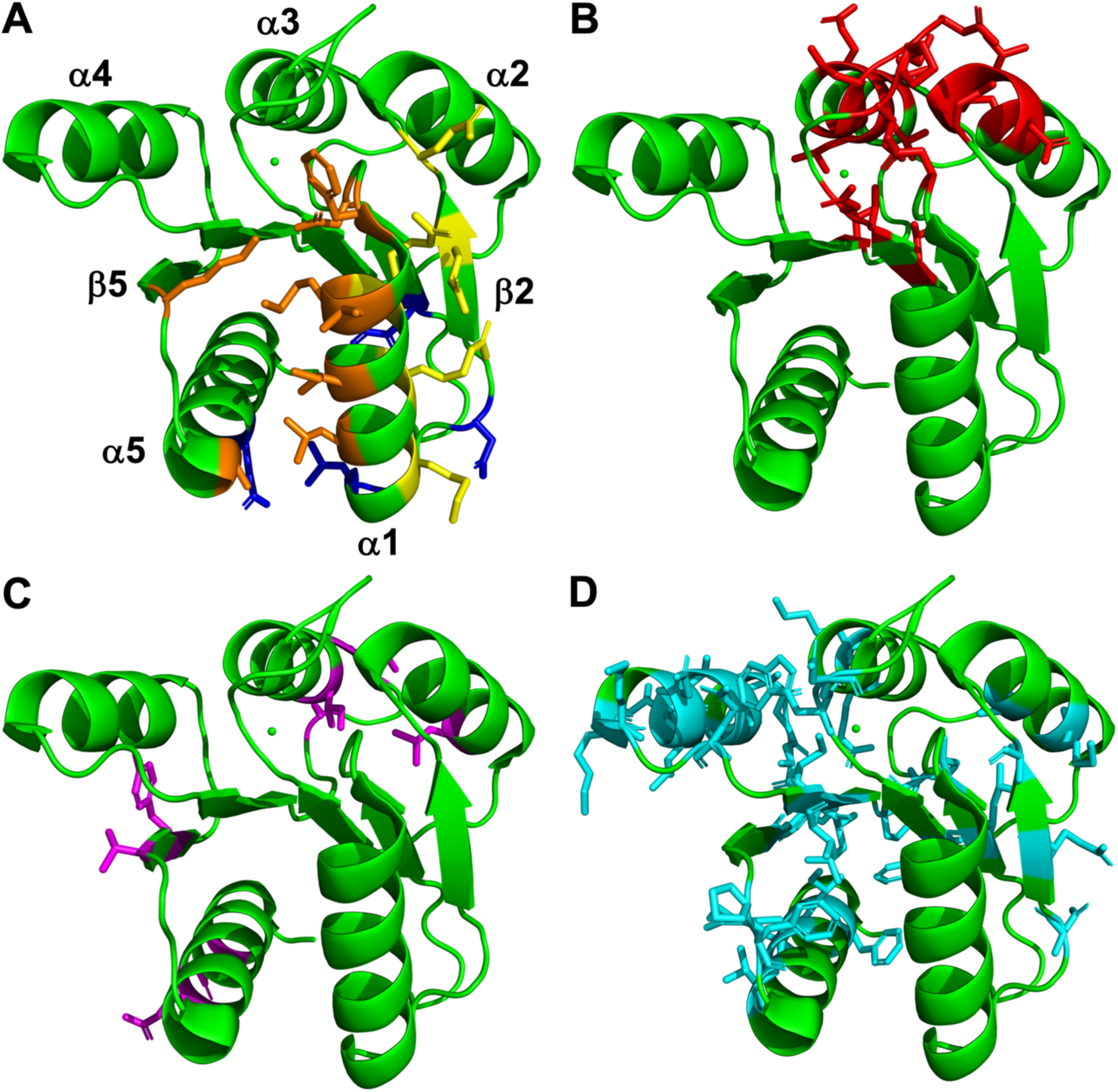
Six inferred networks of interactions within bacterial PsR domains. Positions with the strongest pairwise mutual information scores are mapped onto a receiver domain structure (1fqw). Green dot below α3 is Mg^2+^ ion. Only interactions within the top 2% (core + extended network) are shown. **Table S2** has a complete list of residues for core and extended networks. (A) Networks 1, 2 and 3 (orange, yellow, and blue, respectively) included the catalytic DD1 and K positions conserved in receiver domains, variable positions known to affect receiver domain conformation and/or involve signaling in the α1α5 surface (DD+4, DD+11, K+2, K+4) (47), and positions on the α1β2α2 surface. (B) Network 4 (red) included residues involving the β3α3 loop and surrounding helices, including position D+4, a highly conserved Pro in receiver domains that is important for the γ turn in the region. (C) Network 5 (magenta) included buried residues running from α2, through the α4β5α5 face, to α5. (D) Network 6 (cyan) contained the most extensive number of residues in two hemispheres, bracketing the active site. One hemisphere may involve linking the α1β2α2 region to the active site, while the other appears to encompass the functionally important α4β5α5 surface.

*Network 1.* A small network with five core residues (orange in **Fig. 3A, Table S2A**) predominantly involving the β5α5α1 surface and perhaps related to the previously described allosteric network connecting D to the α1α5 surface (**Table S1E**). Network 1 was remarkable for including two residues (DD1, K) that are conserved (and therefore do not covary) in receiver domains. Structural and amino acid composition analyses previously led to the hypothesis that functional interactions between the DD1 and K positions are important for PsR function (16).

*Network 2.* A small network with four core residues (yellow in **Fig. 3A, Table S2B**). on the α1β2α2 surface. Although the interaction between core residues DD+5 and D-22 on α1 and β2 was the single strongest mutual information link in both receivers and PsRs, it is unclear what role the α1β2α2 surface plays in either domain. **Figure 2** suggests that the amino acid composition of core positions in Network 2 were similar in receiver and PsR domains. Three of the six possible pairwise combinations between Network 2 core positions exhibited mutual information scores in the top 0.25% of all interactions in PsRs. As described later, several PsRs regulate activity by binding ligands in the vicinity of Network 2.

*Network 3.* A small network of three core residues (blue in **Fig. 3A, Table S2C**) on the opposite side of the PsR domain from D.

*Network 4.* A large network of 16 core residues (red in **Fig. 3B, Table S2D**) centered on the β3α3 loop, which in true receiver domains includes a hairpin γ-turn (**Fig. 1B**).

Network 4 includes numerous buried residues on the α2 and α3 helices. Three of the core residues were among the top 10 positions in PsRs based on cumulative mutual information scores, each participating in 13-15 additional significant interactions. The highly connected residues clustered near the N-terminus of α2 and the β3α3 loop, adjacent to the phosphorylation site in receiver domains. The same region is highly variable in available PsR domain structures (described later and in **Table S3**), suggesting functional importance.

*Network 5.* A moderately sized, long-range network of eight core residues (magenta in **Fig. 3C, Table S2E**) running from the C-termini of α2 and α3 through β5 to exposed residues on α5.

*Network 6.* The most extensive network detected contained 28 core residues (cyan in **Fig. 3D, Table S2F**) including many positions (D+1, D+2, T+2, K+1, K+2) important for phosphorylation-mediated effects in receivers. Network 6 included half of the top 1% of covariation scores observed in PsRs. Altogether, Network 6 included more than one third of all positions retained in the PsR sequence alignment and thus undoubtedly contributes to PsR structure. Six of the top 10 cumulative mutual information scores in PsRs belonged to Network 6 core residues, each participating in 14-23 significant additional pairwise interactions. Indeed, core residues of Network 6 make one to four connections with core residues in each of Networks 1-5. Residues in the Network 6 core seem to fall into two regions, bracketing position D. Positions DD-4 through D-12 formed one hemisphere involving β1, β2, and α2 that may be related to Network 2, which seemingly linked the α1β2α2 surface to the active site. There are two strong covariation connections between Networks 2 and 6 via core residues on β2, suggesting this hemisphere of Network 6 may contribute to the ligand binding role postulated for Network 2. Positions D+16 through K+15 formed a second hemisphere of Network 6 involving β4, α4, β5, and α5. The α4β5α5 surface is functionally important in receiver domains and is the dimerization surface for many PsRs (**Table S3**). Dimerization without phosphorylation appears to be an activation mechanism for many PsRs, as discussed later.

In conclusion, Networks 1 and 6 suggest that regions important for receiver function (e.g. involving the α4β5α5 and β5α5α1 surfaces) are modified in PsR domains to provide function without phosphorylation. In contrast, Networks 2, 3, 4 and 5 suggest that regions (e.g., α1β2α2 surface and α2/β3α3 loop/α3) not recognized as functionally important in receiver domains (4, 8) have assumed functional significance in PsRs

We next sought to examine covariation in PsRs of positions important for allosteric signaling in receiver domains. Direct comparisons between mutual information scores are challenging, because the analysis is heavily dependent on the sequence alignment used. Therefore, we sought to identify interactions that showed the largest absolute rank shifts between receiver domains and PsRs (by subtracting internal ranks based on mutual information scores for both). Because the sequence alignment used for PsRs was initially constructed with receiver domains, the positions used in the covariation analysis were directly comparable. **Fig. 4** reveals the pairwise interactions that exhibited the largest shifts in mutual information “rank” (and therefore changes in putative importance). The positions most commonly represented among mutual information scores exhibiting the largest differences between receivers and PsRs were DD+4 (α1), D+2 (β2α2 loop), D+11 (α3), T+2 (β4α4 loop), T+7 (α4), K-6 (α4β5 loop), K-5 (α4β5 loop), and K+14 (α5). Three quarters of the largest rank shifts involved interactions that were statistically significant in receivers but not in PsRs (red in **Fig. 4**). A noteworthy example involves the interaction between positions DD+4 and T+2, which was in the top 0.5% of all significant shifts. DD+4 and T+2 are in close proximity to each other and the receiver domain active site. T+2 is part of the aforementioned “Y-T” coupling mechanism of allostery that connects the active site to the α4β5α5 surface in response to phosphorylation (43). DD+4 sits at the periphery of the allosteric pathway linking the active site to the α1α5 surface (44). We speculate the change in linkage may reflect the lack of reliance on phosphorylation that these two allosterically important regions exhibit in PsRs as compared to true receiver domains.

**FIG 4.**
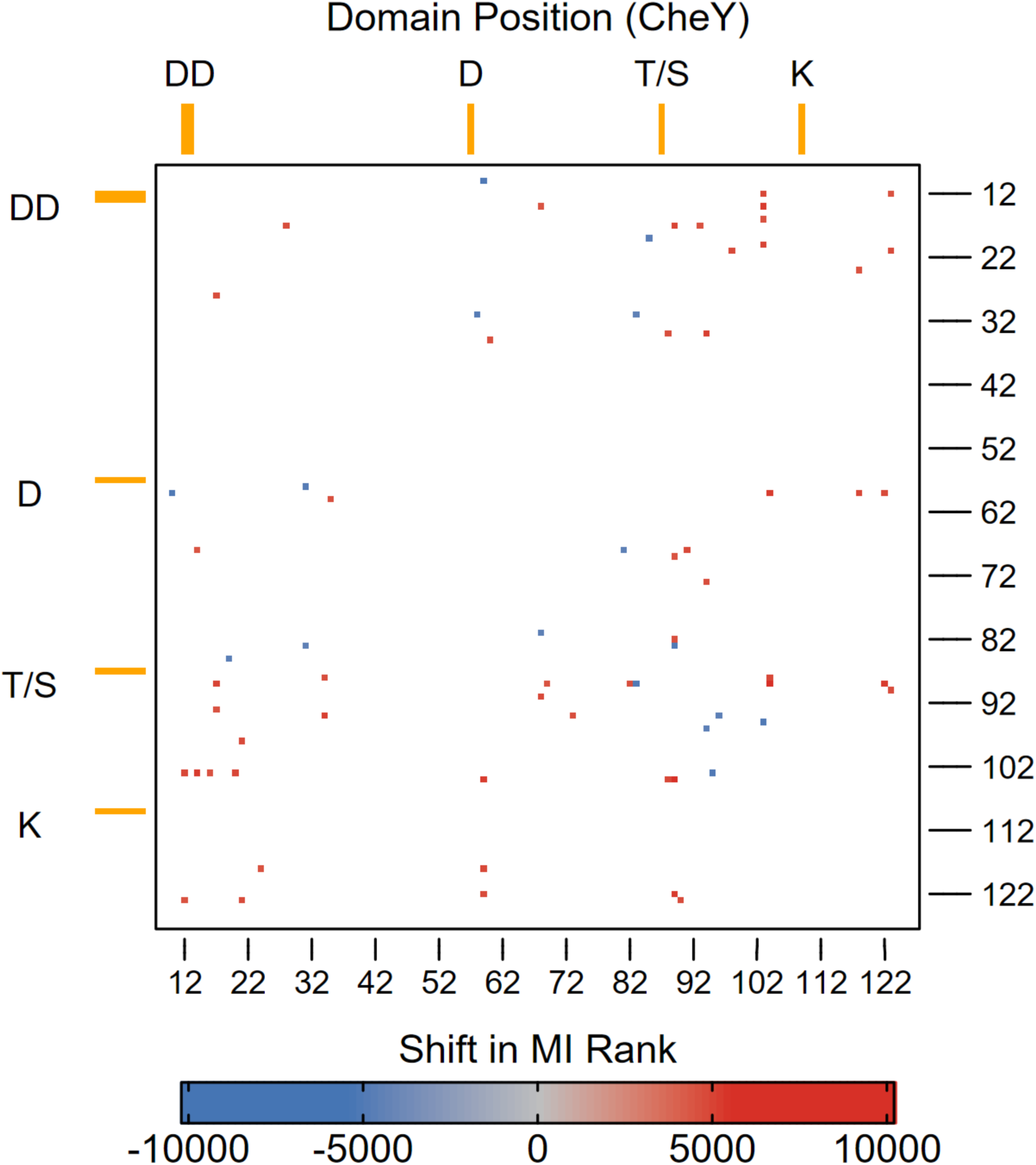
Heatmap of mutual information score rank shifts between receiver and PsR domains. Pairwise mutual information scores for bacterial receiver and PsR domains were constructed as described in Supplemental Material and ranked (higher mutual information = higher rank). Rank shifts were calculated by subtracting PsR ranks from the same pairwise interaction in receiver domains. The top 2% most substantial changes (by absolute rank shift) were included in the heatmap, out of 1,709 significant changes if the interaction was considered statistically significant (mutual information z-score ≥ 6.5 in at least one group). Numbering is according to the *E. coli* CheY sequence. Red colored points indicate that the interaction was “stronger” in receiver domains, whereas blue indicates the opposite.

In summary, various analyses of our bacterial PsR multiple sequence alignment all show that PsR domains often lack features associated with phosphorylation-mediated signaling beyond the simple absence of one or more of the five active site residues conserved in receiver domains.

## Structural analysis of PsR domains

We collated a broader set of bacterial PsR structures than has previously been analyzed. **Table S3** lists 50 high resolution X-ray crystallography and NMR structures of 26 bacterial PsR domains. Forty percent of the PsR domains with known structures arose from structural genomics efforts and so lack corresponding genetic or biochemical studies to connect observed structural features to PsR function.

Receiver domains typically form dimers upon phosphorylation. Fifteen of the available PsR domain structures are dimers in the absence of phosphorylation, a potential mechanism of activation discussed further below. Response regulators are traditionally classified based on their output domains, which co-evolve with the receiver domains (50, 51). Major families of response regulators homodimerize on the α4β5α5 (OmpR/PhoB family), α4β5 (NtrC family), or α1α5 (NarL/FixJ family) interfaces, consistent with known allosteric pathways connecting the phosphorylation site to these surfaces (4). All the observed PsR dimerization interfaces involve some portion of α4β5α5, except for CsgD, FleQ, RitR, and VpsT (**Table S3**).

Consistent with our covariation analysis of PsR domains described above, many differences are observed in the α2 helix, γ turn (β3α3 loop), and α3 helix of PsR structures compared to receivers. As noted later, PsRs that bind ligands often use these regions.

## Key questions and potential hypotheses about mechanism(s) of PsR action

Due to their relationship with TCSs, PsRs are generally assumed to participate in signal transduction, and there is experimental evidence to support this conjecture for many ARRs. By definition, signal transduction requires conversion of an environmental stimulus into an internal representation that is used to implement an appropriate response to changing conditions. In turn, the need to synchronize response with stimulus implies an ability to turn the signal transduction pathway off as well as on. If signal transduction is a primary role for PsRs, then the central unanswered question about PsRs becomes what are the mechanism(s) that regulate PsR activity, i.e., turn PsRs “on’ or “off’?” Theoretically, regulation of activity could be indirect or direct. Each possibility is considered below.

## Indirect mechanisms to regulate constitutively active PsRs

In some models of indirect PsR 2regulation, PsRs are hypothesized to be constitutively active in their ground state (i.e., when synthesized), so the question splits into “What is the source of PsR activity?” and “How are active PsRs neutralized?” Constitutive activity implies that the source of activity is somehow embedded in the PsR primary amino acid sequence.

There are at least three obvious and not mutually exclusive possibilities for how amino acid sequence might lead to constitutive PsR activity:

i. The amino acid at the “D” position. In canonical response regulators, phosphorylation of the D stabilizes active conformations (4). For many response regulators, a longer negatively charged Glu side chain in place of the D can act as a phosphomimetic and partially activate the protein without phosphorylation (52–57), perhaps by binding with the conserved T and K residues to stabilize an active conformation. Therefore, an attractive hypothesis is that the “D” residue of PsRs is the source of constitutive activity. The observation that the most frequent amino acid at the “D” position of ALRs is Glu (16) (**Dataset S3**) is consistent with such a model for many PsRs. However, this category of hypotheses can easily be tested [and has been disproved in some cases (58, 59)] by observing if PsR activity changes upon replacing the “D” residue.
ii. Conformational equilibria. Receiver domains exist in equilibria between conformations that favor or disfavor phosphorylation (mostly the latter). In turn, allosteric pathways reversibly connect the phosphorylation site with the conformations of surfaces involved in mediating responses (4, 44, 60, 61). These connections appear to at least partially form a reciprocal relationship with the catalytic residues. Amino acid composition at key variable positions might bias PsR conformational equilibria toward more “active” configurations. For example, the amino acids at position K+2 strongly affect receiver domain conformation (47). The most abundant residue at K+2 (Phe) biases receiver domains toward inactive conformations, whereas Ala, Ile, Leu, and Val have the strongest effects in shifting receiver domains toward active conformations.

(iii) Multimeric state. Change in multimeric state is a well-known mechanism to regulate protein activity. Many response regulators dimerize upon phosphorylation of their receiver domain to facilitate DNA binding by output domains. Amino acid composition at positions in a multimerization interface might bias PsR equilibria toward multimerization, for example a hydrophobic patch that would be solvent-exposed in monomers but buried in multimers. Consistent with this hypothesis, many ARRs intrinsically form dimers in the absence of phosphorylation, including *Chlamydia trachomatis* ChxR (59, 62, 63), *Pseudomonas aeruginosa* FleQ (64), *Amycolatopsis mediterranei* and *Mycobacterium tuberculosis* GlnR (65), *Helicobacter pylori* HP1043 (66), and *Streptomyces coelicolor* RamR (67). The D residue is important for GlnR and RamR function not via phosphorylation but for dimer formation (65, 67, 68). The *Myxococcus xanthus* ARR FrzS is predicted to form dimers via a C-terminal coiled-coil domain (69).

In contrast to activation, experimental evidence concerning how active PsRs might be neutralized is sparse. Constitutively active PsRs appear to be regulated by spatial or temporal availability, e.g., synthesis, degradation, sequestration, sub-cellular localization, etc., which in turn might be regulated. In *Agrobacterium tumefaciens* and related species, the ARR Rem is inhibited by binding to MirA, whose expression is controlled by a different TCS (58, 70). Expression of *H. pylori* HP1043 is regulated post-transcriptionally or post-translationally, in addition to transcriptionally (71).

## Indirect mechanisms to regulate PsR activity through protein/protein interactions

Another category of indirect regulation involves changes in interaction of PsR domains with other domains or proteins due to stimulus detection through the latter. For example, in *M. xanthus*, binding of GTP to MglA results in binding of MglA•GTP to SgmX, which in turn uncovers a binding site on SgmX for the α3β4α4 surface of the FrzS PsR domain (72). The net result is to bring SgmX to the cell pole where FrzS is located and activate Type IV pili. To the best of our knowledge, there are no reports of receiver domains using the α3β4α4 surface to interact with other proteins. The *Caulobacter crescentus* response regulator PleD has three domains: receiver, PsR, and diguanylate cyclase. Phosphorylation of the receiver domain by partner sensor kinases results in changes in interactions between the receiver and PsR domains, which leads to dimerization and diguanylate cyclase activity (73–75).

The *Synechococcus elongatus* ARR NblR does not require phosphorylation for activity. NblR binds to other proteins in two-hybrid assays (76, 77), but the biological significance of such interactions has apparently not been evaluated. In *Streptomyces venezuelae*, the BldM response regulator does not require phosphorylation for activity (78), is not known to have a partner sensor kinase, is not demonstrably phosphorylated by small molecule phosphodonors, and forms homodimers to regulate gene expression (79). A branched Val residue at position T+1 (39) may sterically hinder phosphorylation of BldM. The ARR WhiI does not form homodimers but regulates a different set of genes by forming heterodimers with BldM The heterodimer is effectively a coincidence detector that activates expression of particular genes only when both constituent monomers are expressed.

A generalization of the WhiI/BldM case is that a PsR could exert its effects by interacting with other components of a TCS. This possibility seems particularly relevant for *Clostridioides difficile* CmrRST, in which the CmrT ARR is part of what appears to be an otherwise normal TCS composed of the CmrS sensor kinase and CmrR response regulator (80). The genes for all three proteins are part of the same operon and therefore presumed to work together somehow. CmrT theoretically might affect CmrS/CmrR phosphoryl group reactions in many ways. For example, CmrT might bind to CmrS to affect autophosphorylation of CmrS or phosphotransfer to CmrR [such binding competitors were hypothesized by Collins & Childers (24)]. Formation of heterodimers between PsR and receiver domains could depend on the phosphorylation state of the receiver. Furthermore, there are several examples of cooperative phosphorylation in response regulators (81–83). The mechanism is believed to be that a phosphorylated monomer stabilizes an active conformation of an unphosphorylated monomer within a heterodimer, leading to enhanced phosphorylation of the second monomer. Similarly, CmrT PsR might promote an active conformation of the CmrR receiver within a heterodimer to promote autophosphorylation of CmrR with small molecule phosphodonors or phosphotransfer from CmrS to CmrR. Finally, CmrT might affect dephosphorylation kinetics of CmrR-P within a heterodimer.

An alternative class of models for integration of a PsR into a TCS is that instead of the PsR interacting with and altering phosphoryl group reactions of the sensor kinase and/or response regulator, the TCS (e.g., CmrRS) might respond to environmental stimuli in the usual way and implement a response simply by changing expression levels of a constitutively active PsR (e.g., CmrT). Consistent with this model, CmrR autoactivates transcription of *cmrRST* and promotes CmrT-dependent phenotypes (84). This strategy would be a simple way to exploit positive autoregulation, which is a feature common to many TCSs (31).

## Direct mechanisms of PsR regulation

There are at least three known mechanisms by which PsR activity can be directly regulated, two of which are variations on receiver domain regulation:

i. Regulated covalent modification. Phosphorylation of the D residue typically regulates receiver but not PsR domains. The *S. coelicolor* ARR GlnR is phosphorylated on multiple Ser/Thr residues and acetylated on multiple Lys residues (85). The covalent modification pattern is dependent upon growth media. Ser/Thr phosphorylation correlates with nitrogen availability and affects GlnR binding to various nitrogen regulatory genes. Lys acetylation also alters DNA binding by GlnR.

Another possibility is that some PsRs that retain the Asp phosphorylation site may function as less efficient versions of response regulators. Amino acid substitutions at DD, T, or K do not necessarily completely ablate phosphorylation and dephosphorylation reactions (86–88).

(ii) Multimeric state. Many response regulators are activated by phosphorylation-mediated dimerization via receiver domains, but PsRs are typically not regulated by Asp phosphorylation. Instead, the *Streptococcus pneumoniae* ARR RitR is activated by dimerization through a disulfide bond between a single Cys residue in the linker between the PsR and DNA-binding domains (89). Dimerization is controlled by redox conditions, primarily the presence of hydrogen peroxide. Because disulfide bond formation and hence dimerization is reversible, RitR effectively acts as a redox sensor. Similarly, the orphan ARRs *H. pylori* Hp1021, *Campylobacter jejuni* Cj1608, and *Arcobacter butzleri* Abu0127 are transcriptional regulators whose activity is redox regulated via dimerization of a Cys at DD+19 in the α1β2 loop (90–93). Multimers of FleQ and VpsT ARRs change conformation dramatically upon c-di-GMP binding at sites distant from the PsR domains. A FleQ4•FleN2 complex formed in the absence of c-di-GMP rearranges to a FleQ3•FleN2 complex in the presence of c-di-GMP (94). For VpsT, the dimer interface between PsR domains changes (95).
(iii) Ligand binding. Receiver domains are not known for binding small molecules other than phosphodonors and metal ions. In contrast, ligand binding to PsRs can alter the activity of the protein in which the PsR domain resides. Five examples follow. In many cases, ligand binding to PsR domains occurs in regions that seem to differ between PsRs and receivers. The PsR domain of the *C. crescentus* hybrid histidine kinase ShkA binds to the HisKA domain and inhibits autophosphorylation (96). Binding of c-di-GMP to the α4β5α5 surface (commonly used in receivers for dimerization and/or downstream partner binding) of the PsR domain stabilizes an alternate conformation in which the HisKA domain has access to the receiver domain, allowing phosphorylation of HisKA and receiver to proceed. Somewhat similarly, the PsR domain of the *S. elongatus* hybrid histidine kinase CikA inhibits autophosphorylation (97). Quinones bind to the α1β2 region of the CikA PsR domain and lead to CikA degradation (98, 99). The α1β2α2 surface of the CikA PsR also binds to KaiB (100). Quinones bind near the β3α3 loop and the N-terminal end of α3 of the PsR domain of *S. elongatus* KaiA and affect the ability of KaiA to stimulate KaiC (101–103). DNA binding by ARRs *S. venezuelae* JadR1 and *S. coelicolor* RedZ are inhibited by binding the end products (jadomycin B and undecylprodigiosin respectively) of the biosynthetic pathways that each control (104).

## Unknown mechanisms of PsR regulation

The activity of the *M. xanthus* ARR FruA is regulated post-translationally by C-signaling during fruiting body formation (105). The mechanism of activation remains unknown, but decades of investigation have effectively ruled out phosphorylation at the conserved Asp. Direct mechanisms such as covalent modification elsewhere or ligand binding are prime candidates.

The ARRs *P. aeruginosa* AtrR (106), *Streptomyces pristinaespiralis* PapR6 (107), and *Streptomyces virginiae* VmsT (108) are functionally important regulators of expression of specific genes in known pathways, but the mechanisms underlying activity of the PsR domains have not been established. Binding of small molecules is proposed to regulate DNA binding by PapR6 analogous to JadR1 and RedZ, but experimental tests did not support the hypothesis. VmsT may be constitutively active, because production of VmsT is regulated by VmsR, a *Streptomyces* Antibiotic Regulatory Protein.

## Future directions in bacterial PsR research

We contend that the molecular mechanisms of activation and inactivation are major unknown aspects of bacterial PsR function. The covariation analyses reported here suggest that focusing experimental investigation on the roles of positions K+1/K+2, as well as the regions encompassed by the α1β2α2 surface, α3, and the β3α3/β4α4 loops is likely to be productive. Progress in determining mechanisms will in turn shed light on the answers to two important questions about bacterial PsRs:

First, what (if any) activation and inactivation mechanisms utilized by bacterial PsRs are shared with canonical receiver domains? For example, positions K+1/K+2 affect receiver domain conformation (47), but β2, α2, the β3α3 loop, and α3 have not been assigned critical roles in receiver domain function. Uncertainty about similarities and differences between molecular mechanisms used by receiver and PsR domains has been a primary source of confusion in the field to date.

Second, are there a small number of activation and inactivation mechanisms shared by most bacterial PsRs (i.e., just a few types of PsRs), or are there a large number of mechanisms (i.e., do most PsRs work differently from one another)? So far, the limited number of established mechanisms seem to be diverse. If there are many different mechanisms, then it could be challenging to ascertain general principles governing most PsRs. Response regulators are often classified based on their output domains (4).

Amino acid sequences of PF00072-type ARRs are phylogenetically intermingled with typical response regulators in all major response regulator families, whereas sequences of PsRs from minor domain families are phylogenetically distinct (30). These data suggest PsRs evolved independently many different times. However, it is not clear if functional convergent evolution has occurred. One approach to address the question of PsR diversity could be to perform a cluster analysis (109) of PsR sequences to estimate how many different classes of PsRs appear to exist and determine which sequence features distinguish the different classes, essentially replicating the above analysis at a higher resolution.

## Supporting information

Dataset S3 PsR Amino Acid Frequencies

Dataset S4 True Receiver Amino Acid Frequencies

Dataset S5 Amino Acid Coabundance Calculator for PsRs

Dataset S1 PsR Alignment

Dataset S2 True Receiver Alignment

## ACKNOWLEDGEMENTS

This work was funded by National Institutes of Health grants R01 AI143638 to RT and R01 GM050860 to RBB, as well as bridge funding from the UNC School of Medicine to RBB.

## Notes

### Competing Interest Statement

The authors have declared no competing interest.

