## Supplementary material for "A Clarifying Perspective on Bacterial Pseudo-Receiver Domains": Materials & Methods, Tables S1-S3

For submission to the *Journal of Bacteriology*
6/23/25

### SUPPLEMENTAL MATERIAL

#### A Clarifying Perspective on Bacterial Pseudo-Receiver Domains

Robert B. Bourret[1#], Emily N. Kennedy[1], Rita Tamayo[1], Clay A. Foster[2]

[1]Department of Microbiology and Immunology, University of North Carolina, Chapel Hill, North Carolina, USA

[2]Department of Pediatrics, Section Hematology/Oncology, University of Oklahoma Health Sciences Center, Oklahoma City, Oklahoma, USA

Running Head: Pseudo-Receiver Domains

#### MATERIALS AND METHODS

**Obtaining sequence sets.** UniProtKB (1) entries with corresponding InterPro domains were downloaded from the InterPro database (2) (August 14th, 2023). Entries were filtered to retain only proteins with either a detectible receiver (IPR001789/PF00072; note that this includes PsR domains), Hpt (IPR008207/PF01627), HisKA (IPR003661/PF00512) or HisKA_3 (IPR011712 /PF07730) domain. A list of representative proteomes (3) (specifically RP75) was obtained from the Protein Information Resource database (4). UniProtKB entries described above were further filtered to retain proteins from representative proteomes only (numerous proteomes contained multiple distinct proteins with full receiver domains). Full-length sequences were obtained for the retained entries and initial domain sizes were estimated using the existing InterPro domain boundaries. Taxonomic lineage information was incorporated based on the NCBI Taxonomy Database (5, 6). To minimize taxonomic bias, sequences from genera containing multiple distinct species were subsetted to retain only a single randomly selected representative species. AlphaFold models were used for manual validation and for secondary structure identification. Sequences lacking a corresponding model in the AlphaFold database (v.4) (7-9) were excluded. Finally, sequence sets were filtered to exclude non-bacterial proteins.

**Processing and distinguishing receiver and PsR domain sequences.** A small percentage of bacterial receiver domain sequences exhibited abnormally short domain sizes (< 60 residues), likely indicating incomplete sequencing data, and were excluded. The full-length, single-domain receiver protein *E. coli* CheY was added to the beginning of the set as a reference sequence. To alleviate domain prediction ambiguity at the termini of the sequences, predicted receiver domain boundaries were expanded by 20 residues on both the N- and C-termini (or as much as possible if already near the true terminus of the protein) for the retained sequence set. If the resulting expanded domain length was still < 110 residues (the estimated average receiver domain size), then additional positions were added symmetrically (where possible) until the 110-residue threshold was reached. Sequences shorter than 110 residues after this extension process were excluded, and highly redundant expanded sequences were filtered out using CD-HIT (v.4.8.1; cutoff of 90%) (10). AlphaFold models for the remaining domain sequences were downloaded and secondary structure predictions were extracted from the resulting .cif/.pdb files (based on a three-state model) with the *mkdssp* program (11, 12). More exact domain boundaries were established by identifying subsequences capturing the canonical $(\beta\alpha)_5$ secondary structure topology. If multiple subsequence orientations ("matches") were found, the "correct" one was determined by maximizing the number of catalytic landmarks detected. The secondary structure predictions were used to search for these catalytic landmark residues based on the following criteria: a metal-binding acidic residue pair (DD, ED, EE, or DE; termed DD) near the C-terminus of β1, a phosphorylatable Asp residue (termed D) near the C-terminus of β3, a Ser or Thr (termed T) near the C-terminus of β4, and a Lys (termed K) near the C-terminus of β5. If no ideal $(\beta\alpha)_5$ match was identified for a given sequence, isolated or likely erroneous secondary structure assignments (e.g., two-residue long helices or strands, short coils breaking up otherwise contiguous structures, etc.) were corrected for and the process was repeated. Domains lacking $(\beta\alpha)_5$ topology even after these corrections were excluded from the data set. Extraneous flanking residues outside of the predicted central $(\beta\alpha)_5$ topology of a canonical receiver domain were trimmed from each sequence. Additionally, some receiver domains feature a highly extended α5 helix (often connecting to an output domain). Because this might throw off alignment of the core conserved domain structure, these extended helices were truncated to a maximum of 17 residues (the overall average length of α5 in the sequence set). A given sequence was classified as a PsR if it lacked any of the four catalytic landmarks mentioned above. Of note, many PsRs encode a Glu in place of (or sometimes in addition to) the phosphorylatable Asp. In this scenario, the presence of the D landmark was confirmed if an Asp residue was closer to the C-terminus of β3 than any additional acidic residue. The above protocol resulted in 9,153 PsR sequences (out of a total of 152,269 sequences) or ~6% of our non-redundant representative bacterial sequence set. WITCH (WeIghTed Consensus Hmm alignment; witch-msa v.1.0.7) (13) was utilized to generate a sequence alignment of all retained sequences using default parameters and then split into separate PsR (**Dataset S1**) and receiver domain (**Dataset S2**) sub-alignments. The two sub-alignments were analyzed to determine pairwise mutual information scores and amino acid abundances at key positions in the domain relative to the *E. coli* CheY reference sequence. To improve interpretability, gaps in the alignments were deleted in relation to the CheY sequence in both sub-alignments.

**Amino acid composition and mutual information analysis.** The bacterial PsR and true receiver domain multiple sequence alignments obtained as described above were submitted to the MISTIC server for corrected mutual information analysis (14). Amino acid frequency tables (**Datasets S3 and S4**) were manually calculated using the seqinr package (v.4.2-36) (15) and converted to percentages. A difference matrix was created to demonstrate shifts in amino acid residue frequencies by subtracting the frequencies detected in the PsR alignment from those found in the true receiver domain alignment. Therefore, positive values represent higher frequency in bona fide receiver domains, whereas negative values represent higher frequency in PsRs.

#### DATASETS

**Dataset S1. Multiple sequence alignment of 9,153 bacterial PsR sequences in FASTA format**

*E. coli* CheY added as a reference. Available as a separate file.

**Dataset S2. Multiple sequence alignment of 143,116 bacterial true receiver domain sequences in FASTA format**

Already includes *E. coli* CheY. Available as a separate file.

**Dataset S3. Amino acid frequency by position in bacterial PsR sequences.**

Calculated from Dataset S1 as described in Materials and Methods. Available as a separate file.

**Dataset S4. Amino acid frequency by position in bacterial true receiver domain sequences.**

Calculated from Dataset S2 as described in Materials and Methods. Available as a separate file.

**Dataset S5. Amino acid co-abundance calculator for bacterial PsRs.**

Based on Dataset S1. Calculates the number of amino acids at any PsR position *j* given a specific amino acid at position *i*. The multiple sequence alignment has 120 positions corresponding to positions 8-127 of the representative receiver domain *E. coli* CheY. Note that positions are not specified as 8 through 127, but rather as D ± n, where D corresponds to position 57 (the conserved phosphorylation site of receiver domains). Similarly, position DD1 is D-45, DD2 is D-44, T is D+30, and K is D+52. Inclusion of *E. coli* CheY in calculations for simplicity introduces a negligible ~0.01% error. Available as a separate file.

#### TABLES

**Table S1. Amino acid frequencies at key receiver and PsR domain positions**

| Position | Amino Acid | Amino Acid Composition (%)[a] Rec | PsR | Function |
|---|---|---|---|---|
| *A. Active Site Positions* | | | | |
| DD1 | Asp | 62 | 58 | Indirect metal binding |
| | Glu | 38 | 24 | |
| DD2 | Asp | 100 | 28 | Direct metal binding |
| | Asn | 0 | 19 | |
| | Ser | 0 | 9 | |
| | Pro | 0 | 8 | |
| | Ala | 0 | 7 | |
| | Gly | 0 | 7 | |
| D | Asp | 100 | 63 | Phosphorylation site |
| | Glu | 0 | 12 | |
| T | Thr | 69 | 38 | Coordinate phosphoryl group |
| | Ser | 31 | 32 | |
| | Ala | 0 | 11 | |
| K | Lys | 100 | 64 | Coordinate phosphoryl group |
| | Arg | 0 | 11 | |
| *B. Catalytic Variable Positions* | | | | |
| DD+1 | Glu | 30 | 11 | Metal binding? |
| | Asp | 29 | 32 | |
| | Asn | 11 | 14 | |
| | His | 10 | 7 | |
| | Ser | 9 | 12 | |
| D+2 | Met | 20 | 4 | Interaction with attacking or leaving group |
| | Arg | 15 | 9 | |
| | Asn | 12 | 9 | |
| | Gln | 10 | 7 | |
| T+1 | Ala | 52 | 29 | Access to phosphorylation site |
| | Gly | 22 | 14 | |
| | Ser | 10 | 14 | |
| | Thr | 8 | 5 | |
| T+2 | Tyr | 17 | 5 | Interaction with attacking or leaving group |
| | Arg | 14 | 8 | |
| | His | 11 | 6 | |
| | Lys | 10 | 4 | |
| | Phe | 10 | 4 | |

*C. Structural Positions*

| | | | | |
|---|---|---|---|---|
| D+4 | Pro | 88 | 53 | Hairpin turn in β3α3 loop |
| D+8 | Gly | 97 | 73 | N-terminus of α3 helix |

*D. Allosteric Pathway – D to α4β5α5 Surface*

| | | | |
|---|---|---|---|
| D+1 | Ile | 34 | 13 |
| | Leu | 25 | 25 |
| | Val | 22 | 14 |
| T-2 | Met | 23 | 15 |
| | Phe | 18 | 11 |
| | Val | 18 | 21 |
| | Ile | 16 | 14 |
| | Leu | 13 | 23 |
| | Ala | 11 | 9 |
| T | Thr | 69 | 38 |
| | Ser | 31 | 32 |
| | Ala | 0 | 11 |
| T+2 | Tyr | 17 | 5 |
| | Arg | 14 | 8 |
| | His | 11 | 6 |
| | Lys | 10 | 4 |
| | Phe | 10 | 4 |
| K-3 | Tyr | 62 | 43 |
| | Phe | 24 | 22 |
| | Val | 4 | 11 |

*E. Allosteric Pathway – D to α1α5 Surface*

| | | | |
|---|---|---|---|
| DD+4 | Ile | 23 | 12 |
| | Val | 17 | 13 |
| | Leu | 15 | 11 |
| | Asn | 12 | 8 |
| DD+8 | Leu | 48 | 46 |
| | Ile | 17 | 17 |
| | Val | 13 | 9 |
| DD+11 | Leu | 22 | 25 |
| | Ile | 12 | 10 |
| | Ala | 11 | 10 |
| DD+12 | Leu | 78 | 77 |
| T-3 | Ile | 48 | 41 |
| | Leu | 24 | 23 |
| | Val | 22 | 24 |

| | | | |
|---|---|---|---|
| T-1 | Leu | 50 | 34 |
| | Val | 16 | 9 |
| | Ile | 14 | 19 |
| K+1 | Pro | 80 | 71 |
| K+2 | Phe | 40 | 20 |
| | Val | 15 | 20 |
| | Ile | 12 | 13 |
| | Leu | 6 | 16 |
| | Ala | 6 | 6 |
| K+4 | Pro | 23 | 27 |
| | Leu | 12 | 8 |
| | Ala | 8 | 11 |
| K+7 | Leu | 72 | 69 |
| | Ile | 8 | 8 |
| | Val | 8 | 9 |

[a] From Datasets S3 and S4.

**Table S2. PsR domain networks inferred from mutual information analysis**

| Position | *E. coli* CheY Position | Location[a] | Inter-network interactions (position) | Comments |
|---|---|---|---|---|
| *A. Network 1 (orange in Fig. 3A)* | | | | |
| Core network[b] | | | | |
| DD+1 | 14 | N-terminal α1 | | |
| DD+3 | 16 | N-terminal α1 | | |
| DD+4 | 17 | α1 | | |
| DD+7 | 20 | α1 | 6 core (105) | |
| K | 109 | β5α5 loop | | |
| Extended network[b] | | | | |
| DD1 | 12 | β1α1 loop | | |
| DD+11 | 24 | α1 | | |
| K+4 | 113 | N-terminal α5 | | |
| *B. Network 2 (yellow in Fig. 3A)* | | | | |
| Core network | | | | |
| DD+5 | 18 | α1 | | In top 10 highest PsR MI[c] scores |
| DD+11 | 22 | α1 | | |
| D-22 | 35 | β2 | 6 core (34; 36) | In top 10 highest PsR MI scores |
| D-20 | 37 | β2α2 loop | 4 core (41) | |
| Extended network | | | | |
| DD+13 | 26 | C-terminal α1 | | |
| *C. Network 3 (blue in Fig. 3A)* | | | | |
| Core network | | | | |
| DD+15 | 28 | C-terminal α1 | 4 core (83); 6 core (58; 59) | In top 10 highest PsR cMI[d] scores |
| DD+18 | 31 | α1β2 loop | | |
| K+8 | 117 | α5 | | |
| Extended network | | | | |
| DD-5 | 8 | N-terminal β1 | | |

*D. Network 4 (red in Fig. 3B)*

Core network

| | | | | |
|---|---|---|---|---|
| DD2 | 13 | β1α1 loop | 6 core (58; 59) | |
| D-19 | 38 | N-terminal α2 | | |
| D-18 | 39 | N-terminal α2 | 6 core (11) | In top 10 highest PsR MI/cMI scores |
| D-16 | 41 | α2 | 2 core (37) | |
| D-14 | 43 | α2 | | |
| D-1 | 56 | C-terminal β3 | | |
| D+3 | 60 | β3α3 loop | | In top 10 highest PsR MI/cMI scores |
| D+4 | 61 | β3α3 loop | | |
| D+5 | 62 | β3α3 loop | | |
| D+6 | 63 | β3α3 loop | | In top 10 highest PsR MI/cMI scores |
| D+7 | 64 | N-terminal α3 | | |
| D+8 | 65 | N-terminal α3 | | |
| D+11 | 68 | α3 | | |
| D+12 | 69 | α3 | 6 core (73; 85) | |
| T-4 | 83 | N-terminal β4 | 3 core (28); 6 core (58) | |

Extended network

| | | | |
|---|---|---|---|
| D-17 | 40 | N-terminal α2 | |
| D-3 | 54 | β3 | 6 core (81) |
| D | 57 | β3α3 loop | |
| D+10 | 67 | α3 | 6 extended (70) |
| D+14 | 71 | α3 | |

*E. Network 5 (magenta in Fig. 3C)*

Core network

| | | | |
|---|---|---|---|
| D-11 | 46 | α2 | |
| T-15 | 72 | α3 | |
| T-12 | 75 | C-terminal α3 | |
| T-11 | 76 | α3β4 loop | |
| K-3 | 106 | β5 | 6 core (99) |
| K-1 | 108 | C-terminal β5 | |

| | | | |
|---|---|---|---|
| K+9 | 118 | α5 | 6 core (115; 119) |
| K+16 | 125 | α5 | 6 core (104) |

<u>Extended network</u>

| | | |
|---|---|---|
| T-8 | 79 | α3β4 loop |

*F. Network 6 (cyan in Fig. 3D)*

<u>Core network</u>

| | | | | |
|---|---|---|---|---|
| DD-4 | 9 | β1 | | |
| DD-2 | 11 | C-terminal β1 | 4 core (39) | In top 10 highest PsR MI scores |
| D-25 | 32 | α1β2 loop | | In top 10 highest PsR MI scores |
| D-23 | 34 | β2 | 2 core (35) | In top 10 highest PsR MI/cMI scores |
| D-21 | 36 | C-terminal β2 | 2 core (35) | In top 10 highest PsR MI scores |
| D-15 | 42 | α2 | | In top 10 highest PsR MI scores |
| D-12 | 45 | α2 | | In top 10 highest PsR MI scores |
| D+1 | 58 | β3α3 loop | 3 core (28); 4 core (13; 83) | In top 10 highest PsR cMI scores |
| D+2 | 59 | β3α3 loop | 3 core (28); 4 core (13) | |
| D+16 | 73 | C-terminal α3 | 4 core (69) | In top 10 highest PsR MI scores |
| T-6 | 81 | α3β4 loop | 4 core (54) | |
| T-5 | 82 | α3β4 loop | | |
| T-2 | 85 | β4 | 4 core (69) | |
| T-1 | 86 | β4 | | |
| T+2 | 89 | β4α4 loop | | |
| T+7 | 94 | α4 | | |
| K-10 | 99 | α4 | 5 core (106) | |
| K-7 | 102 | α4β5 loop | | |
| K-6 | 103 | α4β5 loop | | |
| K-5 | 104 | α4β5 loop | 5 core (125) | In top 10 highest PsR MI/cMI scores |
| K-4 | 105 | N-terminal β5 | 1 core (20) | In top 10 highest PsR MI/cMI scores |
| K+1 | 110 | β5α5 loop | | In top 10 highest PsR MI/cMI scores |
| K+2 | 111 | β5α5 loop | | In top 10 highest PsR MI scores |
| K+6 | 115 | α5 | 5 core (118) | |
| K+10 | 119 | α5 | 5 core (118) | In top 10 highest PsR MI/cMI scores |

| | | | | |
|---|---|---|---|---|
| K+13 | 122 | $\alpha$5 | | |
| K+14 | 123 | $\alpha$5 | | In top 10 highest PsR MI scores |
| K+15 | 124 | $\alpha$5 | | |
| <u>Extended network</u> | | | | |
| D-4 | 53 | N-terminal $\beta$3 | | |
| D+13 | 70 | $\alpha$3 | 4 extended (67) | |
| T-3 | 84 | $\beta$4 | | |
| T+4 | 91 | N-terminal $\alpha$4 | | |
| T+5 | 92 | N-terminal $\alpha$4 | | |
| T+8 | 95 | $\alpha$4 | | |
| T+9 | 96 | $\alpha$4 | | |
| T+10 | 97 | $\alpha$4 | | |
| T+11 | 98 | $\alpha$4 | | |
| K-8 | 101 | C-terminal $\alpha$4 | | |

[a] Based on the structure of *E. coli* CheY (PDB ID 1fqw). N- or C-terminal indicates within three residues of end of $\alpha$ helix or within one residue of end of $\beta$ strand.

[b] Core networks are composed of isolated residue groups intra-connected by mutual information scores in the top 0 to 1%. Extended networks are composed of additional residues connected to existing core groups by mutual information scores in the top 1 to 2%. Only first-order neighbors are considered as part of additional networks via extended interactions. Extended networks were identified by merging core isolated networks with mutual information interactions from the top 1 to 2% within Cytoscape (16) and applying a best neighbor filter.

[c] Mutual Information as calculated by MISTIC (14).

[d] Cumulative MI = sum of all statistically significant (≥ 6.5) MI scores involving the indicated residue. High cMI is often associated with functional and/or catalytic residues (14).

**Table S3. Experimentally determined[a] structures of PsR domains[b]**

| PBD ID | Species | Protein[c] | "DD D T K"[d] | Dimer?[e] | Comments | Source |
|---|---|---|---|---|---|---|
| *PF00072 Response regulator receiver domain (2M InterPro entries)* | | | | | | |
| 3luf | *Aeromonas salmonicida* | A4SNL2 | DD D S Q | $\alpha 4\beta 5\alpha 5$ | 1/2 length, Rec and PsR domains | NYSGXRC |
| 3mf4 | *Aeromonas salmonicida* | A4SNL2 | DD D S Q | $\alpha 4\beta 5\alpha 5$ | 1/2 length, Rec and PsR domains | NYSGXRC |
| 4dad | *Burkholderia pseudomallei* | Q63I75 | ED D T W | $\alpha 4\beta 5$ | $\alpha 2$ kinked, no $\gamma$ turn | JCSG |
| 4dn6 | *Burkholderia pseudomallei* | Q2T2Y8 | ED D S W | $\alpha 4\beta 5$ | $\alpha 2$ kinked, no $\gamma$ turn | JCSG |
| 1w25 | *Caulobacter crescentus* | PleD | DD N V R | $\beta 5\alpha 5$ | Full length PleD•c-di-GMP, Rec (D1) & PsR (D2) domains. "Dimer" interface between Rec and PsR of same chain | (17) |
| 2v0n | *Caulobacter crescentus* | PleD | DD N V R | $\alpha 4\beta 5\alpha 5$ | Full length PleD•$BeF_3^-$•c-di-GMP•GTP$\alpha$S, Rec (D1)& PsR (D2) domains. "Dimer" interface between Rec and PsR of same chain | (18) |
| 2wb4 | *Caulobacter crescentus* | PleD | DD N V R | $\alpha 4\beta 5\alpha 5$ | Full length PleD•$BeF_3^-$•c-di-GMP, Rec (D1) & PsR (D2) domains. "Dimer" interface between Rec and PsR of same chain | Wassman *et al.*, unpublished |
| 6qrl | *Caulobacter crescentus* | ShkA | AS D T K | No | ShkA PsR•c-di-GMP, $\alpha 2$ short, no $\alpha 3$, $\alpha 4$ bent, no $\gamma$ turn, c-di-GMP binds to $\alpha 4\beta 5\alpha 5$, ShkA is a HHK | (19) |
| 6qrj | *Caulobacter crescentus* | ShkA | AS D T K | No | Full length ShkA•AMPPNP, PsR and Rec domains, $\alpha 2$ short, no $\alpha 3$, $\alpha 4$ bent, no $\gamma$ turn, ShkA is a HHK | (19) |
| 2qzj | *Clostridiodes difficile* | CmrT | DG E T K | $\alpha 4\beta 5\alpha 5$ | | NYSGXRC |
| 3lua | *Clostridium thermocellum* | A3DK02 | DY D T K | No | A3DK02 is a HHK | NYSGXRC |
| 2zay | *Desulfuromonas acetoxidans* | Q1JZD9 | DT E S K | $\alpha 4\beta 5\alpha 5$ | | NYSGXRC |

| | | | | | | |
|---|---|---|---|---|---|---|
| 2pln | *Helicobacter pylori* | HP1043 | EK S S K | $\alpha4\beta5\alpha5$ | No $\gamma$ turn | (20) |
| 2hqo | *Helicobacter pylori* | HP1043 | EK S S K | $\alpha4\beta5\alpha5$ | NMR, no $\gamma$ turn, | (20) |
| 2hqr | *Helicobacter pylori* | HP1043 | EK S S K | $\alpha4\beta5\alpha5$ | NMR, full length, no $\gamma$ turn | (20) |
| 2gkg | *Myxococcus xanthus* | FrzS | ES A G K | No | | (21) |
| 2i6f | *Myxococcus xanthus* | FrzS | ES A G K | No | | (21) |
| 2nt3 | *Myxococcus xanthus* | FrzS | ES A G K | No | Y102A mutant | (21) |
| 2nt4 | *Myxococcus xanthus* | FrzS | ES A G K | No | H92F mutant | (21) |
| 3nhm | *Myxococcus xanthus* | Q1CZZ7 | EN D S K | No | No $\alpha4$ | NYSGXRC |
| 3n53 | *Pelobacter carbinolicus* | Q3A6W4 | DQ D S K | No | No $\alpha4$ | NYSGXRC |
| 3kto | *Pseudoalteromonas atlantica* | A0A9J9BB43 | DH E A K | No | | NYSGXRC |
| 3hv2 | *Pseudomonas fluorescens* | Q4K707 | DS A T K | $\alpha4\beta5$ | | NYSGXRC |
| 2j48 | *Synechococcus elongatus* | CikA | EE A L K | No | NMR, $\alpha4$ and $\beta5$ short, CikA is a HHK, Quinones bind to $\alpha1\beta2$ | (22) |
| 5jyu | *Thermosynechococcus elongatus* | CikA | ED E S L | No | NMR, $\alpha2$ short, no $\alpha4$, CikA is a HHK | (23) |
| 5jyv | *Thermosynechococcus elongatus* | CikA | ED E S L | No | NMR, CikA PsR•KaiB, $\alpha2$ short, no $\alpha4$, binds to KaiB on $\alpha1\beta2\alpha2$ surface, CikA is a HHK | (23) |
| 4lzl | *Streptococcus pneumoniae* | RitR | EK N D K | No | $\alpha4$ kinked | (24) |
| 5u8m | *Streptococcus pneumoniae* | RitR | EK N D K | Cys128 | Full length, $\alpha4$ kinked | (25) |
| 5u8k | *Streptococcus pneumoniae* | RitR | EK N D K | No | Full length C128S mutant, $\alpha4$ kinked | (25) |
| 5vfa | *Streptococcus pneumoniae* | RitR | EK N D K | No | Full length C128D mutant, $\alpha4$ kinked | (25) |
| *PF06490 Flagellar regulatory protein FleQ (3K InterPro entries)* | | | | | | |

| | | | | | | |
|---|---|---|---|---|---|---|
| 4wxm | *Pseudomonas aeruginosa* | FleQ | DD G G M | α1 | α2 short, α4 short | (26) |
| 8p53 | *Pseudomonas aeruginosa* | FleQ | DD G G M | α1 | Cryo-EM, full length FleQ•FleN, α2 short, α4 short | (27) |
| 8pb9 | *Pseudomonas aeruginosa* | FleQ | DD G G M | α1 | Cryo-EM, 2/3 FleQ (PsR-AAA+)•c-di-GMP•FleN, α2 short, α4 short | (27) |
| 8h5v | *Vibrio cholerae* | FlrA | ED G N F | No | α2 short, no α4 | (28) |
| *PF21155 VpsT-like, receiver domain (2K InterPro entries)* | | | | | | |
| 3kln | *Vibrio cholerae* | VpsT | SD D C D | α1 | Full length, α2 kinked, α4 short | (29) |
| 3klo | *Vibrio cholerae* | VpsT | SD D C D | α6 | Full length VpsT•di-c-GMP, α2 kinked, α4 short | (29) |
| 5xp0 | *Escherichia coli* | CsgD | TK D T M | α1 | No α2, α4 short | (30) |
| *PF21194 TadZ-like, receiver domain (241 InterPro entries)* | | | | | | |
| 3fkq | *Eubacterium rectale* | TadZ | DK E T K | NA | Full length TadZ•ATP, α3 is a short $3_{10}$ helix, no α4 | (31) |
| *PF21695 Transcription regulator GlnR, N-terminal domain (5K InterPro entries)* | | | | | | |
| 4o1h | *Amycolatopsis mediterranei* | GlnR | TA D V L | α4β5α5 | α1 and α4 kinked, α2 short, no γ turn | (32) |
| 4o1i | *Mycobacterium tuberculosis* | GlnR | TS D V L | α4β5α5 | α1 and α4 kinked, α2 short, no γ turn | (32) |
| 8hih | *Mycobacterium tuberculosis* | GlnR | TS D V L | No | Cryo-EM, GlnR•RNA polymerase•DNA complex, α1 and α4 kinked, α2 short | (33) |
| *PF21714 Circadian clock protein KaiA, N-terminal domain (508 InterPro entries)* | | | | | | |
| 1m2e/ 1m2f | *Synechococcus elongatus* | KaiA | ES V G I | No | NMR, no α4, no γ turn Family of 25 NMR structures | (34) |
| 1r8j | *Synechococcus elongatus* | KaiA | ES V G I | NA | Full length, no α4, no γ turn | (35) |

| | | | | | | |
|---|---|---|---|---|---|---|
| 4g86 | *Synechococcus elongatus* | KaiA | ES V G I | NA | Full length KaiA•DBMIB, no $\alpha 4$, no $\gamma$ turn DBMIB quinone binds near the $\beta 3\alpha 3$ loop/ N-terminal end of $\alpha 3$ | (36) |
| 5c5e | *Synechococcus elongatus* | KaiA | ES V G I | NA | Full length KaiA•KaiC peptide, no $\alpha 4$, no $\gamma$ turn | (37) |
| *<u>PF22368 Response regulator protein ChxR, N-terminal domain (19 InterPro entries)</u>* | | | | | | |
| 3q7r | *Chlamydia trachomatis* | ChxR | EH E D R | $\alpha 4\beta 5\alpha 5$ | No $\alpha 2$ or $\alpha 3$, no $\gamma$ turn | (38) |
| 3q7s | *Chlamydia trachomatis* | ChxR | EH E D R | $\alpha 4\beta 5\alpha 5$ | No $\alpha 2$ or $\alpha 3$, no $\gamma$ turn | (38) |
| 3q7t | *Chlamydia trachomatis* | ChxR | EH E D R | $\alpha 4\beta 5\alpha 5$ | No $\alpha 2$ or $\alpha 3$, no $\gamma$ turn | (38) |
| *<u>No Pfam domain identified</u>* | | | | | | |
| 3snk | *Mesorhizobium loti* | Q989D4 | SS D S K | No | $\alpha 2$ short, no $\gamma$ turn | JCSG |

[a]X-ray unless otherwise noted.

[b]Structure of PsR domain only unless otherwise noted

[c]UNIPROT identifier if protein unnamed

[d]PsR residues corresponding to receiver domain DD D T K conserved residues.

[e]PsR dimer interface indicated if dimer observed. NA (not applicable) indicates dimer but not via PsR domain.

**Structures excluded from Table S3**

- *Pseudomonas aeruginosa* AmiR (PBD ID 1qo0) was identified as containing a PsR domain in 1999 (39). However, this identification has not stood the test of time. The order of secondary structural elements encoded in receiver domain primary sequence is $\beta$1-$\alpha$1-$\beta$2-$\alpha$2-$\beta$3-$\alpha$3-$\beta$4-$\alpha$4-$\beta$5-$\alpha$5 (40), whereas in AmiR the order is $\alpha$1-$\beta$1-$\alpha$2-$\beta$2-$\beta$3-$\alpha$3-

β4-α4-β5-α5 (PBD ID 1qo0). Pfam classifies the domain as PF21332 AmiR, N-terminal, not PF00072 or any of the minor Pfam groupings included in Table S3.

- *Alkalihalobacillus halodurans* BH3024 (PBD ID 2b4a) was identified as a PsR in (24), which is something of a judgement call. BH3024 clearly contains the conserved DD, T, and K residues. Therefore, categorization of BH3024 depends on which residue is identified as corresponding to the D position. The BH3024 residue at the C-terminal end of β3 is a Ser, which presumably is the basis of classification as a PsR. However, the next residue is an Asp, and there are many receiver domains (including the well characterized model receiver domain *E. coli* CheY) in which the phosphorylation site Asp is immediately preceded by a Ser. For the purposes of this review, we regard BH3024 to be a receiver, not a PsR. A definitive experimental test to distinguish between PsR and receiver would be to determine if BH3024 can be phosphorylated on the Asp.
